# Quantification and site-specific analysis of co-occupied N- and O-glycopeptides

**DOI:** 10.1101/2024.07.06.602348

**Authors:** Joann Chongsaritsinsuk, Valentina Rangel-Angarita, Keira E. Mahoney, Taryn M. Lucas, Olivia M. Enny, Mitchelle Katemauswa, Stacy A. Malaker

**Author notes:** To whom correspondence should be addressed: Stacy A. Malaker **Stacy A. Malaker** – Department of Chemistry, Yale University, New Haven, Connecticut 06511.

## Abstract

Protein glycosylation is a complex post-translational modification that is generally classified as N- or O-linked. Site-specific analysis of glycopeptides is accomplished with a variety of fragmentation methods, depending on the type of glycosylation being investigated and the instrumentation available. For instance, collisional dissociation methods are frequently used for N-glycoproteomic analysis with the assumption that one N-sequon exists per tryptic peptide. Alternatively, electron-based methods are indispensable for O-glycosite localization. However, the presence of simultaneously N- and O-glycosylated peptides could suggest the necessity of electron-based fragmentation methods for N-glycoproteomics, which is not commonly performed. Thus, we quantified the prevalence of N- and O-glycopeptides in mucins and other glycoproteins. A much higher frequency of co-occupancy within mucins was detected whereas only a negligible occurrence occurred within non-mucin glycoproteins. This was demonstrated from analyses of recombinant and/or purified proteins, as well as more complex samples. Where co-occupancy occurred, O-glycosites were frequently localized to the Ser/Thr within the N-sequon. Additionally, we found that O-glycans in close proximity to the occupied Asn were predominantly unelaborated core 1 structures, while those further away were more extended. Overall, we demonstrate electron-based methods are required for robust site-specific analysis of mucins, wherein co-occupancy is more prevalent. Conversely, collisional methods are generally sufficient for analyses of other types of glycoproteins.

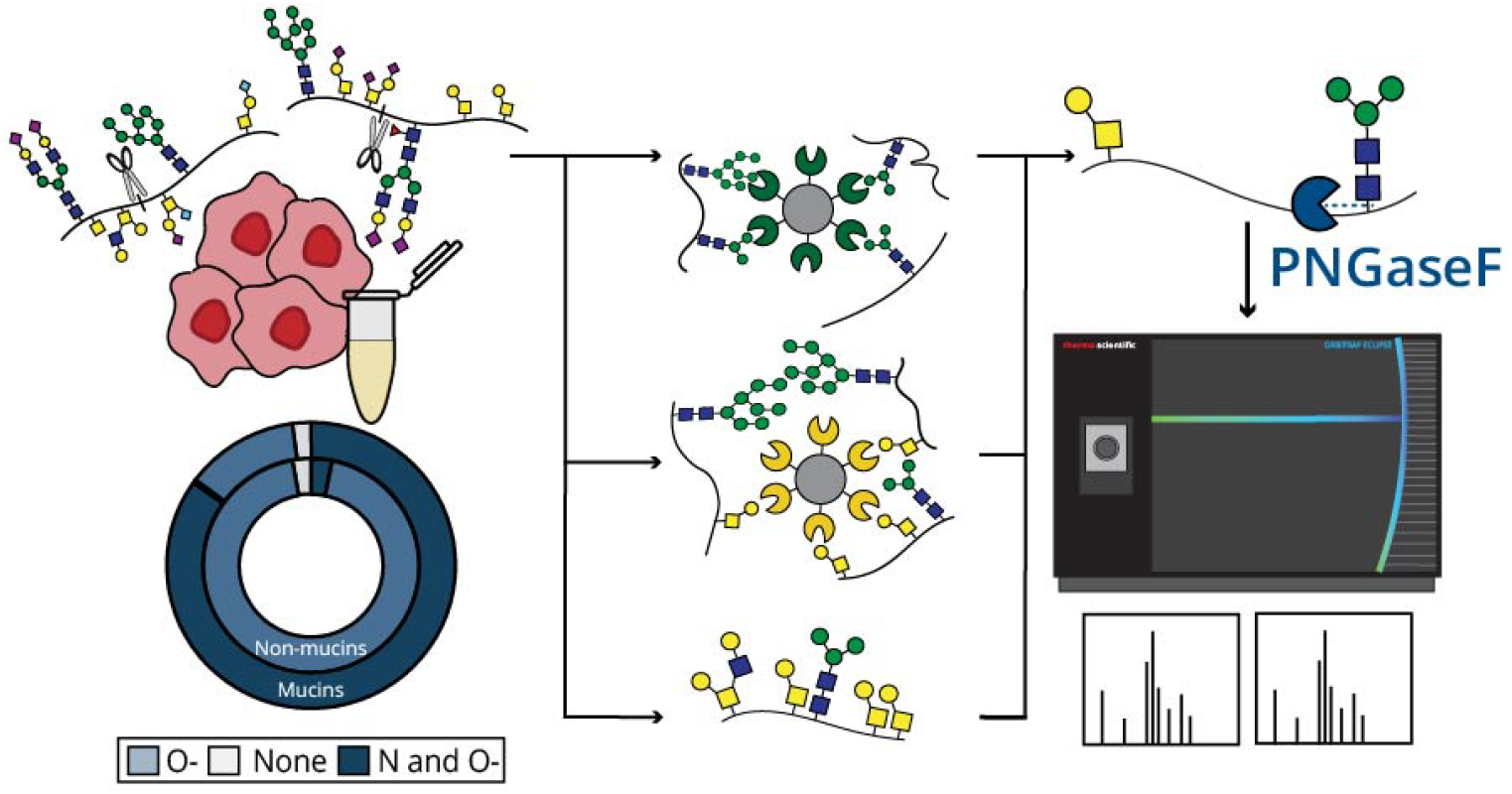

## Introduction

Post-translational modifications (PTMs) are covalent additions of functional groups to proteins which generate large proteomic diversity. PTMs regulate protein activity, localization, turnover, and their interactions, among other important biological processes.^1^ One of the most complex PTMs is glycosylation, whereby proteins are covalently modified with monosaccharides or glycans.^2^ While there are various classes of glycosylation, extracellular protein glycosylation is typically classified as mucin-type O-glycosylation and N-glycosylation.^3^

Mucin-type O-glycosylation (hereafter simply referred to as O-glycosylation) is characterized by an initiating α-N-acetylgalactosamine (GalNAc) attached to the hydroxyl group of any Ser or Thr.^4^ Heterogeneous O-glycan structures are formed through the elaboration of this GalNAc in a non-templated manner. While O-glycans can sparsely decorate proteins, mucins are characterized by dense O-glycosylation within mucin domains, which contain a high density of Pro, Thr, and Ser.^5^ Consequently, mucins exert important biochemical and biophysical functions including cell-cell signaling, barrier formation, and modulation of immune responses.^6-8^ Despite their importance in biological systems, mucins remain understudied since they are difficult to analyze, especially by typical proteomic workflows. For instance, they are resistant to enzymatic digestion by workhorse proteases. To address this issue, O-glycoproteases have recently been introduced and selectively cleave at or near O-glycosylated residues. A subset of O-glycoproteases, called mucinases, are specific for the dense O-glycosylation and bottlebrush-like secondary structure of mucins.^9-13^ Both O-glycoproteases and mucinases can generate O-glycopeptides amenable for MS analysis, thus allowing for improved glycoproteomic sequencing of mucins. That said, the resultant O-glycopeptides often have multiple possible glycosites, necessitating electron-based (ECD or ETD) or hybrid (EThcD or HCD-pd-ETD) fragmentation methods for site-specific glycan localization.^14-16^

N-glycosylation, on the other hand, occurs on Asn residues within an Asn-X-Ser/Thr consensus sequon, where X can be any amino acid except Pro.^17^ N-glycans are characterized by a chitobiose core comprised of three mannose (Man) and two β-N-acetyglucosamine (GlcNAc) residues.^18^ This core structure is further elaborated to form diverse N-glycans of three conventional classes: oligomannose, hybrid, and complex. N-glycans regulate important biological functions including protein folding and stability, solubility, and cell signaling.^19,20^ To simplify glycoproteomic analyses, peptide-N-glycosidase F (PNGaseF) can be used to release most N-glycans by cleaving between the Asn residue and the first GlcNAc of the chitobiose core. PNGaseF converts previously glycosylated Asn residues to Asp via deamidation, a modification that results in a mass difference of +0.98 Da.^21^ Deamidation can also occur stochastically, thus other endoglycosidases which leave ‘scars’ can be used for unambiguous N-glycosite localization. Endoglycosidase H (EndoH) acts on hybrid N-glycans while endoglycosidase F2 (EndoF2) cleaves complex biantennary oligosaccharides and core-fucosylated N-glycans. Both enzymes act on oligomannose N-glycans and, unlike PNGaseF, cleave after the first GlcNAc in the chitobiose core.^22^ Although unambiguous N-glycosite localization is possible with electron-based fragmentation, most tryptic N-glycopeptides contain one N-sequon thus the total glycan composition identified by search algorithms is attributed to the Asn residue. As such, collisional dissociation (CAD or HCD) is considered sufficient for MS analysis within the N-glycoproteomics field, compared to more time-consuming electron-based methods.^23^ Based on a brief literature survey from the last decade, we conservatively estimate that 80% of studies employed collision-based methods for N-glycoproteomic analysis.^24-45^

Despite the standard use of collisional methods for N-glycoproteomics, many groups have reported co-occupied N- and O-glycopeptides from various proteins. For example, in 1990, Escribano *et al*. analyzed alpha-1-microglobulin (AMBP) by RP-HPLC and colorimetric analysis of the hydrolyzed glycans, which implied the presence of both O-linked and N-linked glycosylation.^46^ In another case, Chandrasekhar *et al*. detected N- and O-glycosylation within an N-sequon of potassium voltage-gated channel subfamily E regulatory subunit 1 (KCNE1) by Western Blot analyses of deglycosylated KCNE1.^47^ Mutation of this O-glycosite (T7I) is linked to Jervell and Lange-Nielsen syndrome.^47^ Interestingly, the simultaneous presence of N- and O-glycosylation has been demonstrated to regulate proteolysis. For example, soluble forms of IL-6R (sIL-6R), the receptor for the inflammatory cytokine interleukin-6, are predominantly generated through ADAM-17-mediated proteolysis which is dependent upon glycosylation of both Asn350 and Thr352.^48^ Co-occupancy has also been identified in the spike protein of SARS-CoV-2, where Tian *et al*. observed O-glycosylation of most Ser/Thr within N-sequons, which they referred to as the ‘O-follows-N rule’.^44^ More recently, N- and O-glycopeptides were reported from receptor-type protein tyrosine phosphatase alpha (PTPRA), a regulator of cell adhesion signaling pathways, and corticosteroid-binding globulin protein (CBG), which transports cortisol to inflamed tissues.^49,50^ Within CBG, Ser338 and Asn347 were found to be simultaneously glycosylated, which reduced proteolysis efficiency and thus cortisol bioavailability.^50^ Our laboratory has also detected several deamidated (i.e., previously N-glycosylated) O-glycopeptides from podocalyxin-like protein 1 (podocalyxin), C1 esterase inhibitor (C1-Inh), T cell immunoglobulin and mucin-domain containing protein-3, and T cell immunoglobulin and mucin-domain containing protein-4 (TIM-4) following PNGaseF treatment, suggesting their N- and O-glycosylation.^51,52^ All previously identified co-occupied glycopeptides in the literature are listed in **Supplemental Table 1**.

The presence of simultaneously modified N- and O-glycopeptides suggests that collisional dissociation methods might not be sufficient for routine N-glycoproteomics. Given that four of the ten proteins discussed above were mucins (PTPRA, podocalyxin, C1-Inh, and TIM-4), we predicted the frequency of co-occupancy would be increased in this class of glycoproteins, which naturally contain a greater number of possible O-glycosites. To investigate this, we employed mucin-domain glycoproteins TIM-4, C1-Inh, podocalyxin, and lysosomal associated membrane protein-4 (LAMP-4). Herein, we proteolytically digested these proteins and removed N-glycans using PNGaseF to simplify O-glycosite analysis. For further confirmation of co-occupancy, proteins were digested with the same proteases (or mucinases, where necessary) and treated with EndoH and EndoF2 (**Figure 1A**). We also used these methods to analyze the prevalence of N- and O-glycopeptides in non-mucin glycoproteins including hemopexin, von Willebrand factor (VWF), serotransferrin, and fetuin. To determine the frequency of this phenomenon in complex samples, we enriched glycopeptides from human serum and cell lysate using various glycan-binding proteins (GBPs). Overall, we studied the prevalence of N- and O-glycosylation and the effect of co-occupancy on optimal fragmentation methods for various types of glycoproteomic samples.

**Figure 1.**
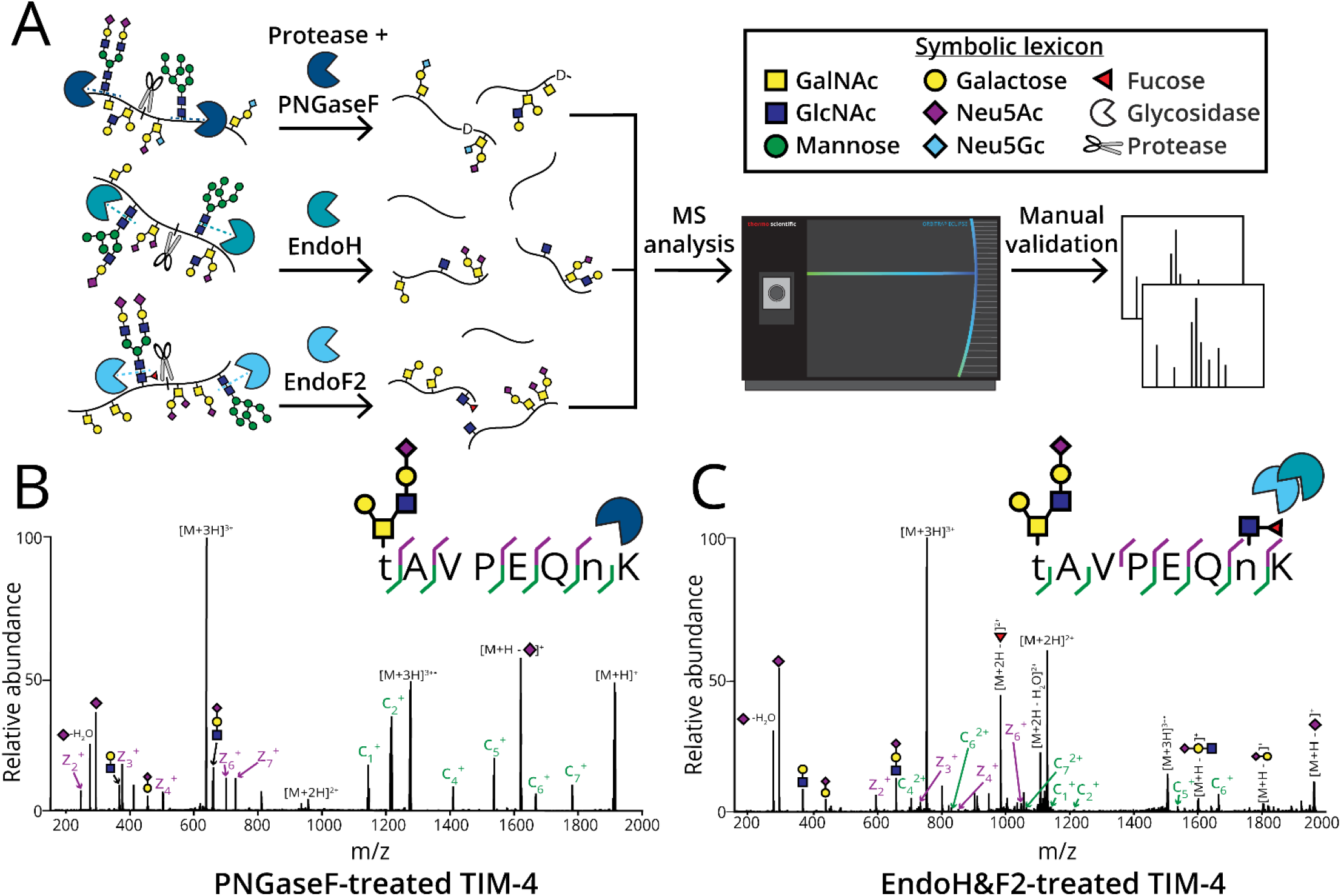
Digestion of glycoproteins for quantification of co-occupancy frequency. A) General workflow for glycoproteomic analysis and quantitation of N- and O-glycopeptides. All proteins were treated with PNGaseF (for quantitation) or EndoH and EndoF2 (to confirm N-glycosite occupancy). The proteins were digested with proteases (or mucinases, where necessary) and the resultant (glyco)peptides were subjected to MS analysis. XICs were generated for each glycopeptide to calculate AUC relative quantitation. B) Recombinant TIM-4 expressed in HEK cells was analyzed with the previously described workflow. ETD enabled site-localization of H2N2A1 to Thr285 in a co-occupied peptide wherein Asn291 was glycosylated, as demonstrated by the spectrum. C) EndoH&F2-treatment of glycoproteins resulted in remaining GlcNAc ‘scars’ for N-glycosite localization of Asn291 and simultaneous O-glycosite localization of H2N2A1 to Thr285 by ETD.

## Results

To quantify the frequency of co-occupied glycopeptides, we digested recombinant or purified mucins and non-mucin glycoproteins with varying combinations of StcE, SmE, and/or IMPa and assessed the resulting (glyco)peptides. Protease conditions were chosen based on protein sequence and are detailed in the experimental section. All cleavage events are shown in **Figure S1**. StcE cleaves within a Ser/Thr-X-Ser/Thr motif, where cleavage occurs N-terminally to a Ser/Thr that is preceded by any amino acid (X) and an O-glycosylated Ser/Thr.^9^ SmE has a less restrictive cleavage motif than StcE and cleaves N-terminally to any O-glycosylated Ser/Thr.^52^ Alternatively, IMPa cleaves N-terminally to any O-glycosylated Ser/Thr, though is restricted by glycosylation on the adjacent P1 position.^12^ Each protein was subsequently digested with trypsin and/or Glu-C and glycosidases overnight. We quantified the presence of N- and O-glycopeptides using PNGaseF-treated samples wherein detected O-glycopeptides were deamidated following N-glycan release (**Figure 1B**). EndoH and EndoF2-treated samples confirmed N-glycosite occupancy through site localization of a GlcNAc or core-fucosylated GlcNAc to an Asn residue with ET(hc)D, as demonstrated in **Figure 1C**. Following MS analysis, RAW files were searched using Byonic. To curate data, (glyco)peptides containing the N-sequon with a score greater than or equal to 200 were manually validated and categorized into four classes: (1) unmodified peptides (those containing the N-sequon wherein the Asn was not deamidated or occupied in Endo-treated data), (2) N-glycopeptides (those that were deamidated following PNGaseF-treatment or were occupied in Endo-treated data), (3) O-glycopeptides (those containing the N-sequon wherein the Asn was not deamidated or occupied in Endo-treated data), and (4) co-occupied N- and O-glycopeptides. We denoted any residues with evidence for glycosylation as lowercase letters. To calculate area-under the curve (AUC) relative quantitation, we extracted ion chromatograms (XICs) of glycopeptides containing the N-sequon. The percent relative abundance of each (glyco)peptide class was calculated from AUCs of PNGaseF-treated samples for determination of co-occupancy frequency.

### Quantification of co-occupancy frequency within mucins

We evaluated the prevalence of N- and O-glycopeptides from biologically relevant mucin proteins TIM-4, C1-Inh, podocalyxin, and LAMP-4. All the proteins studied contained multiple N-sequons except for TIM-4, which only had one potential N-glycosite. Previously, in our characterization of the glycoproteomic landscape of TIM-4, Asn291 was frequently deamidated.^52^ Here, we identified 34 unique O-glycopeptides distributed on 4 peptide backbones from TIM-4 containing the unique Asn291 sequon and confirmed co-occupancy of nearby O-glycosites Thr271, Thr285, and Thr293 (**Supplemental Table 2**). Of these, we found Thr285 was modified 58.7% of the time and Thr293 was modified 39.4% of the time (**Supplemental Table 3**). Overall, the occurrence of co-occupied N- and O-glycopeptides within TIM-4 was 95.9% (**Figure 2A**, navy), while the frequency of O-glycosylation without an N-glycan present was only 4.1% (**Figure 2A**, light blue).

**Figure 2.**
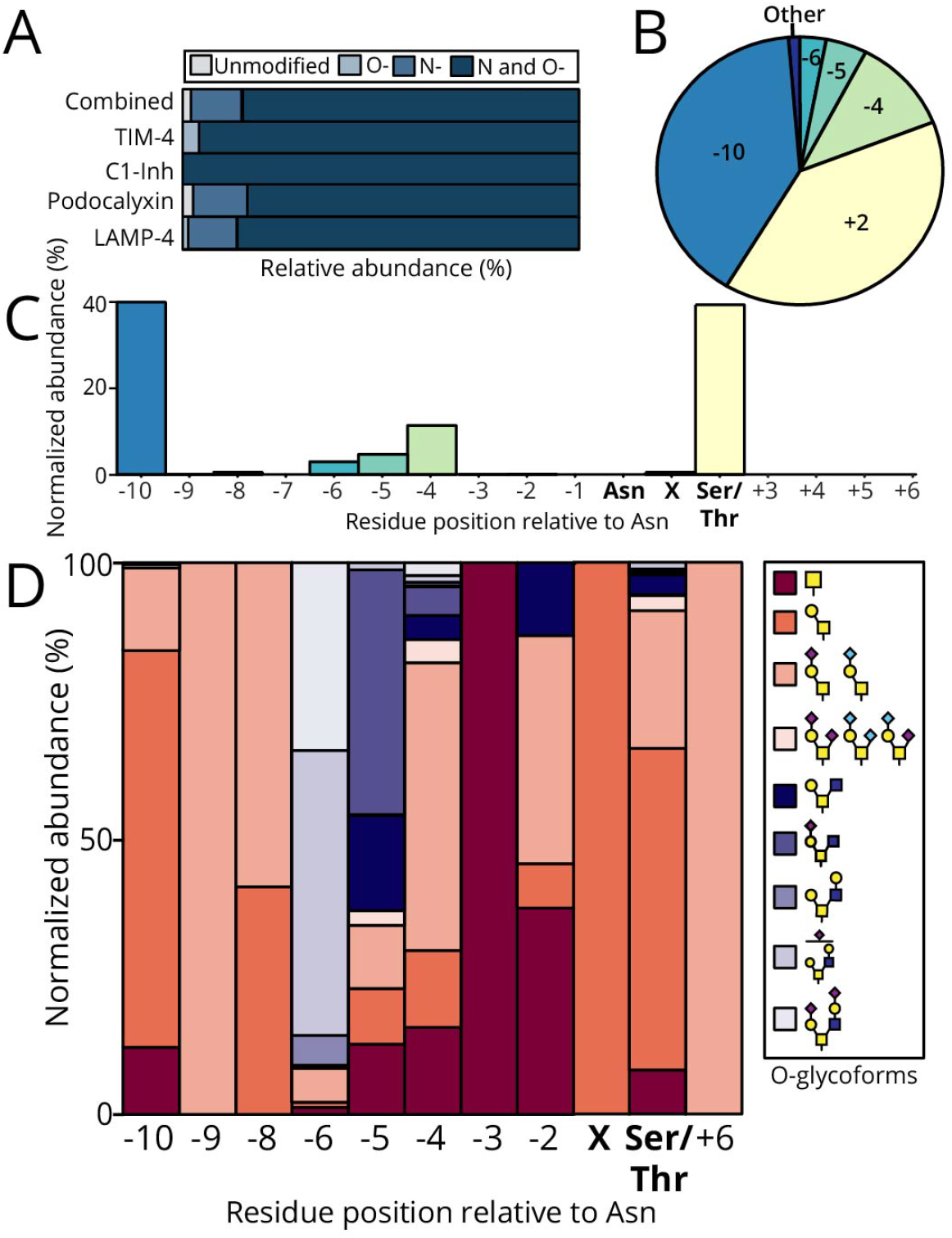
Mucin-domain glycoproteins were frequently co-occupied with predominantly core 1 O-glycan structures. A) Following AUC relative quantitation, percent relative abundances of (glyco)peptide class was calculated for each mucin. The total percent relative abundance was also calculated as an average across all mucins. B) and C) The position of surrounding O-glycosites was normalized to the occupied Asn residue in the N-sequon. Positions N-terminal to the Asn residue were considered negative while those C-terminal to the Asn residue were considered positive. D) The normal relative abundances in percents of glycoforms identified from all mucins was calculated for O-glycosite positions normalized to the occupied Asn. Unelaborated core 1 structures are depicted in a pink gradient while more elongated core 2 structures are indicated by purple hues. X represents the +1 position in the N sequence, while Ser/Thr represents the +2 position.

The next protein we analyzed was C1-Inh, a serine protease inhibitor that is the major regulator of the contact activation, classical, and lectin complement pathways. Previously, 26 O-glycosylation sites were identified on C1-Inh isolated from human plasma, although most were not site-localized.^53^ We also recently characterized the O-glycosylation of C1-Inh with SmE, trypsin, and PNGaseF. In that work, three N-glycosites were deamidated in O-glycopeptides where Thr27, Ser64, Thr71, Thr76, Thr79, and/or Thr83 were glycosylated.^52^ In the current study, we identified 10 unique O-glycopeptides which were all co-occupied (100% frequency; **Figure 2A**) and obtained coverage of two out of three N-sequons within the mucin domain. Specifically, Asn69 was co-occupied with Ser63, Ser64, or Thr67. Furthermore, we also confirmed simultaneous glycosylation of Asn81 with Thr72, Thr76, Thr83, or Thr87 (**Supplemental Table 4**). The most occupied O-glycosite was Thr76 (67.7%), followed by Thr83 (18.3%), and Thr72 (5.2%). All other co-occupied O-glycosites accounted for less than 5% occupancy.

We then examined podocalyxin, a sialoprotein that promotes the growth, proliferation, and metastasis of solid tumors.^54^ The recombinant protein was expressed in NS0 cells originating from mice, which evolutionary retained the gene responsible for production of NeuGc.^55^ As we have previously demonstrated, SmE is not as active on recombinant proteins produced in CHO or NS0 cells.^52^ Thus, we opted to digest podocalyxin with StcE and/or IMPa, along with an additional trypsin digestion, for maximum N-sequon coverage. Podocalyxin contains four potential N-glycosites within its mucin domain: Asn35, Asn45, Asn106, and Asn146. With StcE, we identified Asn106 and Asn146 while with IMPa we were able to detect the other two sites (Asn35 and Asn45). Considering both digestion conditions, we identified 87 unique O-glycopeptides covering all relevant N-sequons. Across these glycopeptides, we unambiguously site-localized Ser23, Ser29, Thr37, Thr40, Thr41, Thr47, Thr95, Thr96, Ser102, Thr107, and Thr108 (**Supplemental Table 5 and 6**). Overall, the total frequency of N- and O-glycopeptides was 78.7% (**Figure 2A**, navy). Lastly, we analyzed LAMP-4, a lysosomal protein containing four potential N-glycosites within its mucin-domain: Asn88, Asn96, Asn118, and Asn126. Of these, we were able to site-localize Asn88, Asn118, Asn126, and 12 O-glycosites, one of which was within the Asn118 sequon. In total, co-occupancy frequency was 80.6% (**Figure 2A**, navy) while the frequency of N-glycosylation without an O-glycan present was 16.2% (**Figure 2A**, medium blue) and conversely, O-glycosylation without an N-glycan present was 0.4% (**Figure 2A**, light blue).

To determine which sites were most frequently O-glycosylated in co-occupied glycopeptides, we normalized their locations in relation to the simultaneously occupied Asn residue, which we considered position zero. Again, we calculated the AUCs which were then normalized to that of the Asn within the N-sequon. Position -10 (light blue) and the Ser/Thr within the N-sequon (light yellow) were both moderately occupied, accounting for 39.6% and 39.0%, respectively (**Figure 2B and 2C**). In contrast, positions -4 (light green), -5 (turquoise), and -6 (teal) were infrequently occupied (11.3%, 4.6%, and 2.9%, respectively). The remaining positions were insignificantly O-glycosylated, with occupancies being less than one percent (**Figure 2B and 2C**). As Tian *et al*. and Chernykh *et al*. suggested that the presence of N-glycosylation sterically hinders the accessibility of O-glycosyltransferases, we were also interested in the O-glycan compositions of the co-occupied glycosites.^44,50^ Less elaborated core 1 structures were depicted in a pink gradient while more elongated core 2 structures were depicted in purple hues (**Figure 2D**). Of note, we looked at glycoform distribution at position -10 and at the Ser/Thr within N-sequons (i.e., +2), which were both moderately occupied. The O-glycosylation at position -10 was not tremendously diverse, where 12.2% was attributed to N1 (maroon) and 15.0% was attributed to H1N1A1/H1N1G1 (peach). The predominant O-glycoform at position -10 (72.1%) was H1N1 (coral), with other glycoforms each accounting for less than one percent abundance (**Figure 2D**). At the Ser/Thr within the N-sequon, H1N1A2/H1N1G2/H1N1A1G1 (light pink) was the least abundant O-glycoform, accounting for 2.7% (**Figure 2D**). N1 and H1N1A1/H1N1G1 were moderately abundant glycoforms (10.5% and 23.2%, respectively) while H1N1 was the predominant glycoform, with an abundance of 54.2% (**Figure 2D**). Generally, we identified core 1 structures at positions proximal to the occupied Asn, with an increased frequency of elongated core 2 structures at sites farther from the occupied Asn (**Figure S2a**). The predominant core 1 structures neighboring the occupied Asn residue were non-sialylated (**Figure S2b**), further emphasizing the lack of their structural complexity. Overall, these findings support the postulation that O-glycans adjacent to occupied N-glycosites tend to be unelaborated due to the steric hindrance provided by N-glycans to the enzymatic machinery responsible for O-glycosylation.

### Quantification of co-occupancy frequency within non-mucin glycoproteins

To investigate whether the frequency of co-occupied peptides were a consequence of prevalent O-glycosylation in mucins, we analyzed serotransferrin, fetuin, hemopexin, and VWF. While these proteins are O-glycosylated, they are not characterized by the dense O-glycosylation characteristic of mucin domains. Serotransferrin, fetuin, and hemopexin contain three, six, and two predicted O-glycosites, respectively.^56^ Indeed, previous O-glycoproteomic analysis of fetuin confirmed occupancy of all six predicted O-glycosites.^57^ Additionally, in-depth O-glycoproteomics of VWF indicated 10 occupied O-glycosites.^36,58^ In this study, we found N-glycopeptides without accompanying O-glycans were the primary species detected across all glycoproteins with frequencies of 99.2%, 99.9%, 99.2%, and 85.2% for serotransferrin, fetuin, hemopexin, and VWF, respectively (**Figure 3A**, cool blue). In contrast to the occurrence of co-occupancy in mucins, a significantly lower frequency (3.2%) of co-occupancy was observed in these glycoproteins (**Figure 3B**, navy). The only protein from which we identified N- and O-glycopeptides was VWF, with a frequency of 8.6% (**Figure 3A**, navy), where Asn2290 and Thr2298 were co-occupied. The predominant O-glycoforms at Thr2298 were sialylated core 1 structures, with di-sialylated core 1 exhibiting a relative abundance of 72.5% (**Figure 3C**, light pink), followed by mono-sialylated core 1 accounting for 21.8% (**Figure 3C**, peach). An example EThcD spectrum of a co-occupied glycopeptide generated from VWF is shown in **Figure 3D**; calculations of co-occupancy for all of these proteins can be found in **Supplemental Table 3**.

**Figure 3.**
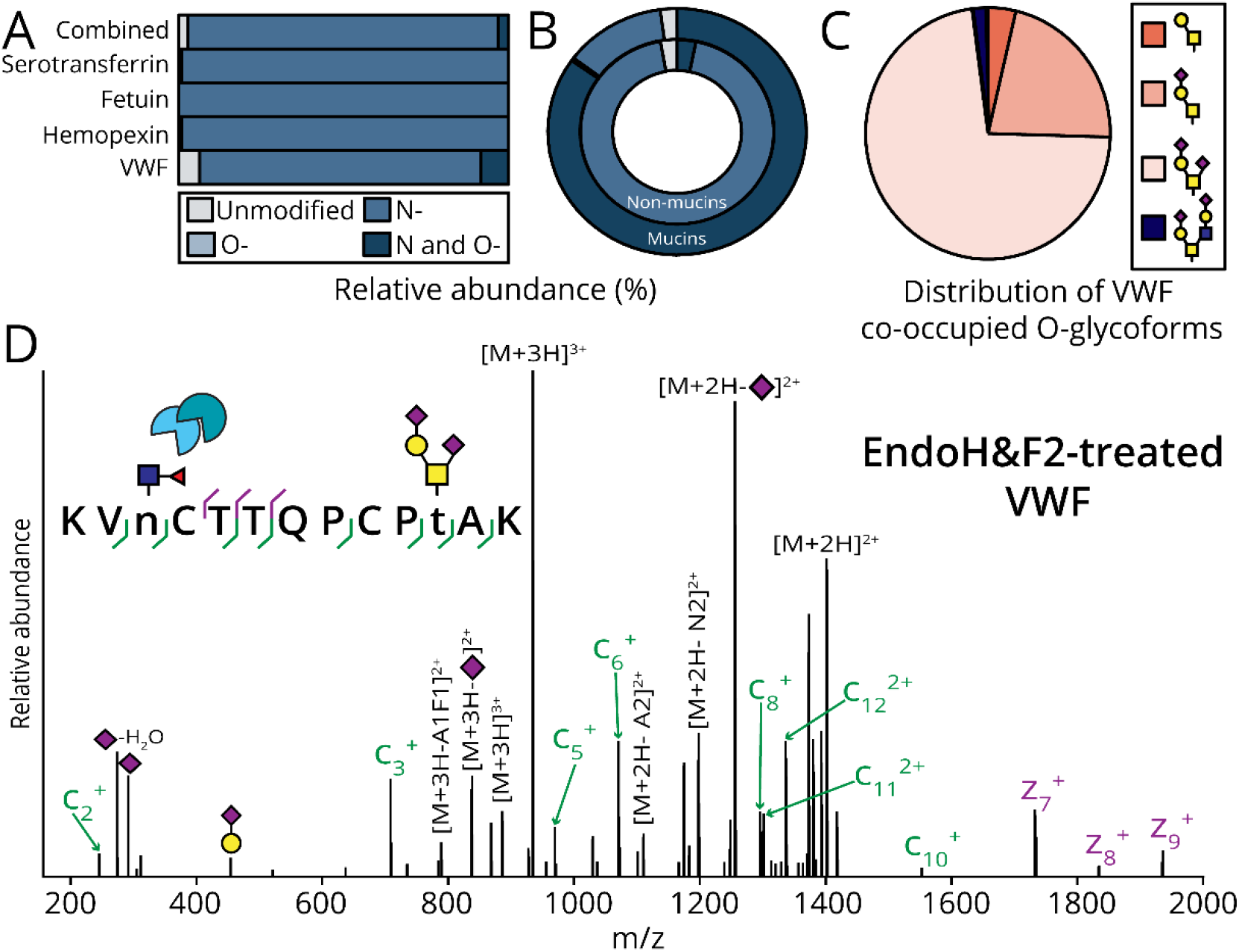
Co-occupancy is an infrequent phenomenon in non-mucin glycoproteins. As described in Figure 2, the percent relative abundances of each (glyco)peptide class were calculated for non-mucin glycoproteins: serotransferrin, fetuin, hemopexin, and VWF. The total percent relative abundance was also calculated as an average across all glycoproteins investigated. B) The relative frequencies of (glyco)peptide classes were compared across non-mucin glycoproteins and mucins in a donut plot. C) The distribution of O-glycan structures decorating Thr2298 was determined following AUC relative quantitation. D) EndoH&F2-treatment of VWF enabled detection of a GlcNAc ‘scar’ by EThcD in LC-MS/MS and consequently N-glycosite localization of Asn 2990 and unambiguous O-glycosite localization of H2N2A2 to Thr2298.

### Quantification of co-occupancy frequency from enrichments of complex samples

To demonstrate that this phenomenon was not an artifact of recombinant protein expression, we investigated the incidence of co-occupancy in complex samples. We enriched for N-glycopeptides, O-glycopeptides, and mucins using a variety of GBPs. As glycopeptides are less abundant than their unmodified counterparts, enrichment is a key step in glycoproteomic analyses of complex samples. We confirmed that most deamidated sites were occupied by N-glycans in our recombinant protein analyses thus we treated all complex sample enrichments with PNGaseF to calculate AUC relative quantitation of co-occupancy.

### N-glycopeptide-enriched HeLa lysate with ConA

Concanavalin A (ConA) is a lectin that binds oligomannose-type N-glycans with high affinity.^59^ To identify co-occupied glycopeptides, we first digested 2 mg of HeLa lysate with SmE and trypsin, followed by a ConA peptide-level enrichment (**Figure 4A**). As we identified a lower prevalence of N- and O-glycosylation from analysis of non-mucin glycoproteins versus mucins, we similarly expected to detect a lower frequency of co-occupied peptides from this N-glycopeptide enrichment. As such, we identified 3374 peptide-spectral matches corresponding to 793 unique deamidated peptides; of these, all were considered to be N-glycopeptides (**Supplemental Table 7, Figure 4B**). This suggested that co-occupied N- and O-glycopeptides are a relatively infrequent occurrence.

**Figure 4.**
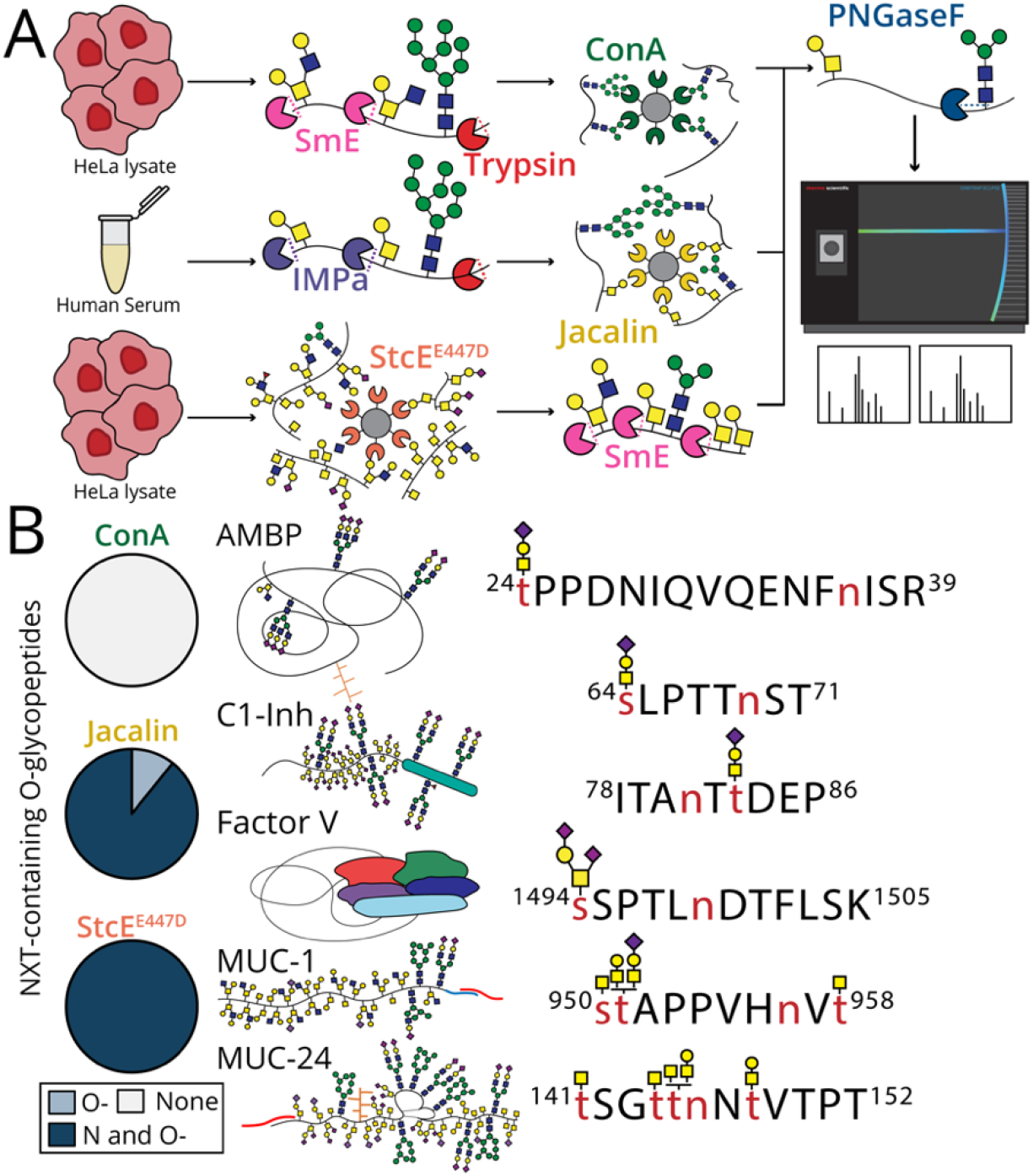
A greater number of co-occupied N- and O-glycopeptides were enriched from mucins in human serum and cell lysate. A) The general workflow for enrichment and glycoproteomic analysis of co-occupied peptides from complex samples is depicted. An N-glycopeptide-level enrichment was performed on HeLa lysate with the lectin, ConA. Similarly, an O-glycopeptide-level enrichment was performed on human serum with the lectin, Jacalin, and mucin-domain glycoprotein-level enrichment was performed on HeLa lysate with StcE^E447D^. All glycopeptides were treated with PNGaseF (for quantitation) and analyzed by LC-MS/MS. XICs were generated for each glycopeptide and AUC quantitation was performed. B) LC-MS/MS analysis was used to identify 14 N- and O-glycopeptides across all enrichments.

### O-glycopeptide-enriched human serum with Jacalin

To enrich for O-glycopeptides from human serum, we used Jacalin, a lectin which has preference for core 1 mucin-type O-glycans.^60^ We digested with IMPa and trypsin then PNGaseF-treated the enriched glycopeptides (**Figure 4A**, yellow). Here, out of 485 glycopeptide-spectral matches (GSMs), five unique N- and O-glycopeptides were identified (**Supplemental Table 8**). We detected two co-occupied glycopeptides from C1-Inh, confirming our analysis of the purified protein. In addition, N- and O-glycopeptides from AMBP, coagulation factor V (FV), and ceruloplasmin were observed. As previously mentioned, AMBP was reported to be modified with both N- and O-glycans following UV-HPLC analysis of hydrolyzed glycans, although site-localization of discrete glycan structures was not possible.^46^ Here, we were able to confirm co-occupancy of Thr24 and Asn36 and site-localized H1N1A1 to Thr24 (**Figure S3**). We also identified glycopeptide s[H1N1A2]SPTLnDTFLSK originating from FV, wherein Ser1494 and Asn1499 were simultaneously occupied (**Supplemental Table 8**). Finally, we detected an N- and O-glycopeptide from ceruloplasmin, EHEGAIYPDn(tt)[H1N1A1]DFQR, which has not been reported before. Although we were unable to site-localize the O-glycan H1N1A1, spectral evidence suggested its presence on either Thr138 or Thr139 (**Supplemental Table 8**). In summary, five unique co-occupied glycopeptides were isolated from a Jacalin enrichment on human serum. These glycopeptides were derived from four proteins, two of which (C1-Inh, CV), are considered mucin glycoproteins.

### Mucin-enriched HeLa lysate with StcE^E447D^

We previously demonstrated that an inactive point mutant of mucinase StcE, called StcE^E447D^, is an effective enrichment tool to probe the mucinome from complex samples including cancer cell lines, human serum, and crude ovarian cancer patient ascites fluid.^61,22^ Here, we performed a protein-level mucin enrichment on HeLa lysate with StcE^E447D^ and digested the enriched mucins with SmE and PNGaseF (**Figure 4A**, orange). In total, 20 GSMs corresponded to seven unique co-occupied peptides (**Supplemental Table 9**). These glycopeptides originated from mucins within the canonical mucin family, including mucin-1 (MUC-1) and sialomucin core protein 24 (MUC-24) (**Figure 4B**). Many canonical mucins are associated with disease progression, especially MUC1, which is aberrantly expressed in ∼60% of all cancers.^63^ Although more is known about MUC-1, MUC-24 is thought to be heavily sialylated with 24 predicted O-glycosites distributed across two mucin domains.^64^ Regardless, the O-glycosites which are occupied and the glycans decorating these sites for both MUC-1 and - 24 still remains to be elucidated. Following our MS analysis, we identified three unique glycopeptides from MUC-1 where glycosylation of Asn957 and Ser950, Thr951, or Thr959 occurred in tandem. In addition, we found co-occupancy of Thr145 or Thr148 with Asn146 across four unique glycopeptides from MUC-24 (**Supplemental Table 9**). A frequency of 100% co-occupancy was detected in the mucin enrichment compared to a frequency of 89.3% co-occupancy in the Jacalin O-glycopeptide enrichment (**Figure 4B**). As in our recombinant protein analysis, this confirmed that mucins exhibit a greater frequency of co-occupancy compared to non-mucin glycoproteins and that a more prevalent synergy exists between N- and O-glycosylation than has been reported.

## Conclusions

Changes in both protein N- and O-glycosylation are considered a hallmark of many diseases. Thus, mapping the glycosylation landscape of proteins at the molecular level holds enormous diagnostic and prognostic value.^65^ Towards this goal, collisional dissociation methods are optimal for N-glycoproteomics. Although electron-based fragmentation is not commonly performed for analysis of N-glycopeptides, numerous studies have reported the prevalence of co-occupied glycopeptides, suggesting this may not be an optimal MS strategy. We sought to determine how frequently co-occupied N- and O-glycopeptides occur and thus determine the need for electron-based methods for analysis in these species.

Here, we demonstrated that the frequency of N- and O-glycopeptides within mucins was greater than in other types of glycoproteins. Accordingly, this work underscored the necessity of both collisional and electron-based methods for comprehensive glycoproteomic analysis of mucins, which is well-acknowledged within the field. Instead, if the goal is to perform routine N-glycoproteomics, collisional dissociation methods are generally sufficient for robust site-specific analysis. Indeed, co-occupancy of only one O-glycosite from non-mucin glycoprotein VWF was observed. Additionally, where N- and O-glycosylation occurred, core 1 structures predominated at O-glycosites proximal to the occupied N-glycosite. This is likely due to the limited accessibility of O-glycosyltransferases following protein exit from the Golgi wherein N-glycosylation takes place, as has been previously postulated.^50^

As our study suggested a higher prevalence of N- and O-glycosylation within mucins, we investigated this frequency on a more global level. Towards this goal, we quantified the occurrence of co-occupancy in enriched glycoconjugates from human serum and HeLa cell lysate. We enriched for N-glycopeptides, O-glycopeptides, and mucins using various GBPs. A total of 13 co-occupied glycopeptides were identified across all enrichments, with a greater number of N- and O-glycopeptides identified from the mucin enrichment. Of these glycopeptides, only two (15.4%) corresponded to non-mucin glycoproteins AMBP and ceruloplasmin while 11 (84.6%) were attributed to mucins.^66^ The high frequency of this phenomenon within mucins suggests a biological relevance for co-occupancy. Of the reported co-occupied glycopeptides to date, all are originated from proteins involved in either transport of biological molecules or cell signaling, indicating that co-occupancy could regulate these biological functions. Overall, our work adds to the ongoing efforts towards the global study of N- and O-glycosylation synergy and its functional roles.

## Materials and methods

### Materials

Recombinantly expressed TIM-4 in HEK293 cells (Q96H15) and podocalyxin-like protein 1 recombinantly expressed in NS0 cells (AAB61574.1) were purchased from R&D Systems (9407-TM, 1658-PD-050). Recombinantly expressed LAMP-4 in HEK293 cells (P34810) was purchased from Elabscience (PKSH034029). C1-Inh (P05155), VWF (P04275), hemopexin (P02790), and human apo-transferrin (P02787) isolated from human plasma were purchased from Sigma Aldrich (E0518), Invitrogen (RP-43132), Athens Research & Technology (16-16-080513), and R&D systems (3188-AT-100MG), respectively. Bovine Fetuin-A (P12763) was purchased from Promega (V4961). The plasmids for His-tagged SmE, StcE, and StcE^E447D^ were kindly provided by the Bertozzi laboratory. SmE, StcE, and StcE^E447D^ were expressed in-house as previously described.^9,52^ The plasmid for PNGaseF expression was purchased from Addgene (Addgene, 114274). PNGaseF was expressed in-house as described and was used for glycopeptide enrichments from complex samples.^67^

### Cell culture

HeLa cells (ATCC, CCL-2) were grown in T175 flasks (Falcon, 353136) and maintained at 37 ºC, 5% CO_2_. The cells were cultured in DMEM (Gibco, 11965-092) supplemented with 10% fetal bovine serum (FBS, Sigma, F0926), 1% sodium pyruvate (Gibco, 11360-070), and 1% penicillin/streptomycin (Gibco, 15140122). The cells were lifted using 1X Hank’s-based enzyme free cell dissociation solution (Sigma, S-004-B).

### Mass spectrometry sample preparation

Unless otherwise mentioned, solutions were made using LCMS grade water (Thermo, T511490-K2), acetonitrile (ACN, Honeywell, LC015), formic acid (Pierce, 85178), and ammonium bicarbonate (AmBic, Honeywell Fluka, 40867).

### Enrichment and digestion of glycopeptides from complex samples

For all enrichments from complex samples, large quantities of PNGaseF were necessary, thus we expressed the enzyme in-house. All samples were desalted prior to MS analysis.

NHS-activated Sepharose 4 Fast Flow beads (Cytiva, 17090601) were washed with 1X PBS (Gibco, 14200-075) three times to remove excess storage solution. Conjugation of 1 mg of StcE^E447D^ or ConA (Vector Laboratories, L-1000) to 1 mL of NHS-activated Sepharose 4 Fast Flow (Cytiva, 17090601) was performed by rotation at 4 ºC for 4 hours. After the reactions, the remaining available binding sites on the beads were blocked twice with 100 mM Tris, pH 7.4 (Sigma, T2663-1L) by rotation at 4 ºC for one hour each. Prior to storage, the beads were rinsed three times with a solution of 20mM Tris, 100mM NaCl (Biotech, SB8889).

For N-glycopeptide enrichment with ConA, 2 mg of HeLa lysate was reduced with a final concentration of 0.5 mM dithiothreitol (DTT, Sigma Aldrich, D0632) shaking at 65 ºC for 30 minutes. Alkylation was performed with a final concentration of 0.75 mM iodoacetamide (IAA, Sigma Aldrich, I1149) at room temperature in the dark for 30 minutes. The sample was digested at 37 ºC with 10 µg of SmE overnight, followed by 10 µL of 0.5mg/mL trypsin (Promega, V511C) for 6 hours. The sample was boiled at 95 ºC for 5 minutes to inactivate the remaining trypsin. N-glycopeptides were enriched by rotation with 250 µL of ConA bead slurry for 6 hours at 4 ºC. To remove non-specific binders and remaining detergent, the beads were washed three times with 1X TBS (Bio-Rad, 1706435), followed by three washes with 20 mM Tris (Sigma, T2663-1L). To elute the enriched glycopeptides, the sample was boiled twice in 250 µL of 0.5% SDC (Research Products International, D91500-25.0) at 95 ºC shaking for 5 minutes each. The sample was digested with 5 µg of PNGaseF at 37 ºC overnight.

The Jacalin peptide level enrichment was performed using 200 µL of Jacalin-agarose bead slurry (Vector Laboratories, AL-1153). Here, 2 mg of pooled human serum (Innovative Research, ISER10mL) was reduced and alkylated as previously described and digested with 5 µL of 0.5mg/mL trypsin at 37 ºC overnight. Prior to rinses and elution, the peptides were boiled at 95 ºC to deactivate any remaining trypsin. The Jacalin bead slurry was washed three times with a solution of 20mM Tris, 100mM NaCl to remove excess storage solution. O-glycopeptides generated from human serum were added to the Jacalin beads and rotated at 4 ºC for 6 hours.

The beads were washed three times with 1X TBS and twice with 20 mM Tris. Finally, the glycopeptides were eluted twice using 200 µL of 0.8 M galactose (Thermo Fisher Scientific, A12813.18) prepared in 20 mM Tris by rotation at 4 ºC for 30 minutes each time. Following elution, samples were desalted to remove the remaining galactose and resuspended in 20 mM Tris. PNGaseF digestion was performed with 3 µg of enzyme overnight at 37 ºC.

Mucin enrichment using StcE^E447D^ was previously described elsewhere.^61,62^ Here, 8 mg of HeLa lysate at a concentration of 2 mg/mL in 10 mM EDTA (Invitrogen, 15575-038) was rotated with 250 µL of StcE^E447D^ bead slurry at 4 ºC for 6 hours. The beads were rinsed and eluted as previously mentioned for the ConA-enrichment, with the addition of 1mM EDTA to reduce enzymatic activity of StcE^E447D^. Reduction and alkylation was performed as previously described and the sample was digested with 8 µg of PNGaseF and 4 µg of SmE at 37 ºC overnight.

### Purified and recombinant protein digestion

Aliquots containing 10 µg of recombinant or purified protein were reconstituted at a concentration of 1 µg/µL in 50 mM AmBic, with 2 µg of protein used for each digest. All proteins were reduced with a final concentration of 2 mM DTT and reacted at 65 ºC for 20 minutes. Samples were then alkylated in 5 mM IAA for 15 minutes in the dark at room temperature. Each protein was digested with PNGaseF (NEB, P0705) or EndoH (NEB, P0702S) and EndoF2 (NEB, P0772S). PNGaseF-treated samples were used for co-occupancy quantitation while EndoH and EndoF2-treated samples were used to confirm N-glycosite occupancy due to stochastic deamidation. For PNGaseF-treated samples, a 1:10 dilution of concentrated PNGaseF was prepared in 50 mM AmBic with a final concentration of 2 mM CaCl_2_ (MP Biomedicals, 153502). Where applicable, mucinase digestion and PNGaseF digestion were performed together at 37 ºC overnight using 1 µL of diluted PNGaseF and StcE, SmE, or IMPa (NEB, P0761). For EndoH and EndoF2-treated samples, 20 µL of 40 mM ammonium acetate (Sigma-Aldrich, A7330) at pH 2-3 was added to each sample to lower the pH to 4-5 for optimal endoglycosidase enzymatic activity. Dilutions of EndoH and EndoF2 at 1:10 were prepared in 20 mM Tris. Samples were incubated with 1 µL of diluted EndoH and EndoF2 at 37 ºC for 8 hours. The pH of the samples was returned to 7 with a final concentration of 2 mM CaCl_2_ and 7 µL of 500 mM AmBic, followed by mucinase digestion. Digestions with SmE and StcE were performed at a 1:10 enzyme to substrate (E:S) ratio while IMPa was used according to manufacturer’s instructions. Digestions were completed with Glu-C and/or trypsin at a 1:40 E:S ratio for 6 hours at 37 ºC.

### Sample desalting

All reactions were quenched with 200 µL of 0.5% formic acid prior to desalting. Desalting was performed using 10 mg Strata-X 33 µm polymeric reversed phase SPE columns (Phenomenex, 8B-S100-AAK). Each column was activated with 1000 µL ACN followed by 1000 µL of 0.1% formic acid, and 1000 µL of 40% ACN, 0.1% formic acid. The column was equilibrated with 1000 µL of 0.1% formic acid. After equilibration, the samples were vortexed, centrifuged, and added to the column. The sample tube was rinsed with 200 µL of 0.1% formic acid, vortexed, and centrifuged then added to the column. The column was additionally rinsed with 200 µL of 0.1% formic acid. The columns were then transferred to a 1.5 mL Eppendorf tube and glycopeptides were eluted by twice with 150 µL of 0.1% formic acid in 40% ACN each time. The eluent was collected into one Eppendorf tube and dried using a vacuum concentrator (LabConco) prior to reconstitution in 10 µL of 0.1% formic acid.

### Mass spectrometry data acquisition

Samples were analyzed by online nanoflow liquid chromatography-tandem mass spectrometry using an Orbitrap Eclipse Tribrid mass spectrometer (Thermo Fisher Scientific) coupled to a Dionex UltiMate 3000 HPLC (Thermo Fisher Scientific). For each analysis, 2 to 4 µL was injected onto an Acclaim PepMap 100 column packed with 2 cm of 5 µm C18 material (Thermo Fisher, 164564) using 0.1% formic acid in water (solvent A). Peptides were then separated on a 15 cm PepMap RSLC EASY-Spray C18 column packed with 2 µm C18 material (Thermo Fisher, ES904) using a gradient from 0-35% solvent B (0.1% formic acid with 80% acetonitrile) in 60 minutes.

Full scan MS1 spectra were collected at a resolution of 60,000, an automatic gain control (AGC) target of 3e5, and a mass range from 300 to 1500 *m/z*. Dynamic exclusion was enabled with a repeat count of 2, repeat duration of 7 s, and exclusion duration of 7 s. Only charge states 2 to 6 were selected for fragmentation. MS2s were generated at top speed for 3 s. HCD was performed on all selected precursor masses with the following parameters: isolation window of 2 *m/z*, 30% normalized collision energy, orbitrap detection (resolution of 7,500), maximum inject time of 50 ms, and a standard AGC target. An additional ETD fragmentation of the same precursor was triggered if 1) 3 of 8 HexNAc or NeuAc fingerprint ions (126.055, 138.055, 144.07, 168.065, 186.076, 204.086, 274.092, and 292.103) were present at ± 0.1 m/z and greater than 5% relative intensity, 2) the precursors were present at a sufficient abundance for localization using ETD. If the precursor was between 300-850 m/z and over 3e4 intensity, an ETD scan was triggered and read out using the ion trap at a normal scan rate. For precursors between 850-1500 m/z with an intensity greater than 1e5, supplemental activation (EThcD) was also applied at 15% nCE and read out in the Orbitrap at 7500 resolution. Both used charge-calibrated ETD reaction times, 150 ms maximum injection time, and 200% standard injection targets.

### Mass spectrometry data analysis

Raw files were searched using Byonic against directed databases containing the relevant protein sequence or proteome. Mass tolerance was set to 10 ppm for MS1’s and 20 ppm for MS2’s. For EndoH and EndoF2-treated samples, carbamidomethyl Cys was set as a fixed modification and N1 and N1F1 were added as rare N-glycan modifications. Asn deamidation was added as a variable modification for PNGaseF-treated samples. For most samples, we used the default O-glycan database containing 9 common structures without H1N1A3 and N2. For analysis of mucin-domain glycoproteins, three glycans were allowed as a common modification while 2 glycans were allowed as a common modification for analysis of glycoproteins. H2N2 was added as an additional O-glycan structure in all searches. For analysis of podocalyxin, NeuGc structures (H1N1G1, H1N1G2, and H1N1A1G1) were included. Files generated using an O-glycoprotease and additional GluC or trypsin digestion were searched with fully specific cleavage N-terminal to Ser and Thr and C-terminal to Arg and Lys and 6 allowed missed cleavages. Samples treated with only GluC and/or trypsin were searched with fully specific cleavage C-terminal to Arg or Lys and 2 missed cleavages. The results were filtered to a score of ≥200. All glycopeptide identities were manually validated using Xcalibur software (Thermo Fisher Scientific). The mass spectrometry proteomics data have been deposited to the ProteomeXchange Consortium via the PRIDE partner repository with the dataset identifier PXD053713.

Reviewer access details:

Project accession: PXD053713

Token: 7ynttXPwu7yS

### Abundance calculations

We extracted XICs of monoisotopic masses and calculated AUCs. To avoid bias toward smaller glycan structures while minimizing the inconvenience of typing multiple isotope masses into XCalibur software (Thermo Fisher Scientific) for each peptide, abundances were calculated using a variable number of isotopes. Only the 12C peak was used for anything under 1600 Da, the 13C was also included up to 2400 Da, and three isotopes were collected over 2400 Da. These values were chosen based on the predicted isotopic distributions for given intact peptide values to allow the majority of the signal to be incorporated. All charge states were included when determining the abundance of a given glycoform. Glycopeptide abundances were calculated using the PNGaseF-treated file.

## Acknowledgements

The authors thank Vincent Chang for expression of StcE^E447D^. J.C. is supported by an NSF GRFP (DGE-2139841); V.R.A. is supported by the Peter Moore Fellowship through the Chemistry Department at Yale University; K.E.M. and T.M.L. are supported by a Yale Endowed Postdoctoral Fellowship in the Biological Sciences; O.M.E. is supported by the National Institutes of Health Biophysical Training Grant (T32GM008283); M.K. is supported by a University Fellowship through Yale University. S.A.M. is supported by NIGMS R35-GM147039.

## References

1. Mann, M., & Jensen, O. N. (2003). Proteomic analysis of post-translational modifications. Nature Biotechnology, 21(3), 255–261. 10.1038/nbt0303-255

2. Bagdonaite, I., Malaker, S. A., Polasky, D. A., Riley, N. M., Schjoldager, K., Vakhrushev, S. Y., Halim, A., Aoki-Kinoshita, K. F., Nesvizhskii, A. I., Bertozzi, C. R., Wandall, H. H., Parker, B. L., Thaysen-Andersen, M., & Scott, N. E. (2022). Glycoproteomics. Nature Reviews Methods Primers, 2(1), 48. 10.1038/s43586-022-00128-4

3. Varki, A., & Kornfeld, S. (2022). Historical Background and Overview. In A. Varki, R. D. Cummings, J. D. Esko, P. Stanley, G. W. Hart, M. Aebi, D. Mohnen, T. Kinoshita, N. H. Packer, J. H. Prestegard, R. L. Schnaar, & P. H. Seeberger (Eds.), Essentials of Glycobiology (4th ed.). Cold Spring Harbor Laboratory Press. http://www.ncbi.nlm.nih.gov/books/NBK579927/

4. Brockhausen, I., Wandall, H. H., Hagen, K. G. T., & Stanley, P. (2022). O-GalNAc Glycans. In A. Varki, R. D. Cummings, J. D. Esko, P. Stanley, G. W. Hart, M. Aebi, D. Mohnen, T. Kinoshita, N. H. Packer, J. H. Prestegard, R. L. Schnaar, & P. H. Seeberger (Eds.), Essentials of Glycobiology (4th ed.). Cold Spring Harbor Laboratory Press. http://www.ncbi.nlm.nih.gov/books/NBK579921/

5. Gendler, S. J., & Spicer, A. P. (1995). Epithelial Mucin Genes. Annual Review of Physiology, 57(1), 607–634. 10.1146/annurev.ph.57.030195.003135

6. Naughton, J., Duggan, G., Bourke, B., & Clyne, M. (2014). Interaction of microbes with mucus and mucins: Recent developments. Gut Microbes, 5(1), 48–52. 10.4161/gmic.26680

7. Van Putten, J. P. M., & Strijbis, K. (2017). Transmembrane Mucins: Signaling Receptors at the Intersection of Inflammation and Cancer. Journal of Innate Immunity, 9(3), 281–299. 10.1159/000453594

8. Kuo, J. C.-H., Gandhi, J. G., Zia, R. N., & Paszek, M. J. (2018). Physical biology of the cancer cell glycocalyx. Nature Physics, 14(7), 658–669. 10.1038/s41567-018-0186-9

9. Malaker, S. A., Pedram, K., Ferracane, M. J., Bensing, B. A., Krishnan, V., Pett, C., Yu, J., Woods, E. C., Kramer, J. R., Westerlind, U., Dorigo, O., & Bertozzi, C. R. (2019). The mucin-selective protease StcE enables molecular and functional analysis of human cancer-associated mucins. Proceedings of the National Academy of Sciences, 116(15), 7278–7287. 10.1073/pnas.1813020116

10. Shon, D. J., Malaker, S. A., Pedram, K., Yang, E., Krishnan, V., Dorigo, O., & Bertozzi, C. R. (2020). An enzymatic toolkit for selective proteolysis, detection, and visualization of mucin-domain glycoproteins. Proceedings of the National Academy of Sciences, 117(35), 21299–21307. 10.1073/pnas.2012196117

11. Rangel-Angarita, V., & Malaker, S. A. (2021). Mucinomics as the Next Frontier of Mass Spectrometry. ACS Chemical Biology, 16(10), 1866–1883. 10.1021/acschembio.1c00384

12. Vainauskas, S., Guntz, H., McLeod, E., McClung, C., Ruse, C., Shi, X., & Taron, C. H. (2022). A Broad-Specificity O -Glycoprotease That Enables Improved Analysis of Glycoproteins and Glycopeptides Containing Intact Complex O -Glycans. Analytical Chemistry, 94(2), 1060–1069. 10.1021/acs.analchem.1c04055

13. Yang, W., Ao, M., Hu, Y., Li, Q.K., & Zhang, H. (2018). Mapping the O-glycoproteome using site-specific extraction of O-linked glycopeptides (EXoO). Molecular Systems Biology, 14(e8486). 10.15252/msb.20188486.

14. Riley, N. M., Malaker, S. A., & Bertozzi, C. R. (2020). Electron-Based Dissociation Is Needed for O-Glycopeptides Derived from OpeRATOR Proteolysis. Analytical Chemistry, 92(22), 14878–14884. 10.1021/acs.analchem.0c02950

15. Khoo, K.-H. (2019). Advances toward mapping the full extent of protein site-specific O-GalNAc glycosylation that better reflects underlying glycomic complexity. Current Opinion in Structural Biology, 56, 146–154. 10.1016/j.sbi.2019.02.007

16. Thaysen-Andersen, M., Wilkinson, B. L., Payne, R. J., & Packer, N. H. (2011). Site-specific characterisation of densely O -glycosylated mucin-type peptides using electron transfer dissociation ESI-MS/MS. ELECTROPHORESIS, 32(24), 3536–3545. 10.1002/elps.201100294

17. Thaysen-Andersen, M., Packer, N. H., & Schulz, B. L. (2016). Maturing Glycoproteomics Technologies Provide Unique Structural Insights into the N-glycoproteome and Its Regulation in Health and Disease. Molecular & Cellular Proteomics, 15(6), 1773–1790. 10.1074/mcp.O115.057638

18. Moremen, K. W., Tiemeyer, M., & Nairn, A. V. (2012). Vertebrate protein glycosylation: Diversity, synthesis and function. Nature Reviews Molecular Cell Biology, 13(7), 448–462. 10.1038/nrm3383

19. Stanley, P., Moremen, K. W., Lewis, N. E., Taniguchi, N., & Aebi, M. (2022). N-Glycans. In A. Varki, R. D. Cummings, J. D. Esko, P. Stanley, G. W. Hart, M. Aebi, D. Mohnen, T. Kinoshita, N. H. Packer, J. H. Prestegard, R. L. Schnaar, & P. H. Seeberger (Eds.), Essentials of Glycobiology (4th ed.). Cold Spring Harbor Laboratory Press. http://www.ncbi.nlm.nih.gov/books/NBK579964/

20. Esmail, S., & Manolson, M. F. (2021). Advances in understanding N-glycosylation structure, function, and regulation in health and disease. European Journal of Cell Biology, 100(7–8), 151186. 10.1016/j.ejcb.2021.151186

21. Tretter, V., Altmann, F., & März, L. (1991). Peptide-N 4 -(N -acetyl-β-glucosaminyl)asparagine amidase F cannot release glycans with fucose attached α1 → 3 to the asparagine-linked N -acetylglucosamine residue. European Journal of Biochemistry, 199(3), 647–652. 10.1111/j.1432-1033.1991.tb16166.x

22. Lee, J. E., Fusco, M. L., & Saphire, E. O. (2009). An efficient platform for screening expression and crystallization of glycoproteins produced in human cells. Nature Protocols, 4(4), 592–604. 10.1038/nprot.2009.29

23. Riley, N. M., Malaker, S. A., Driessen, M. D., & Bertozzi, C. R. (2020). Optimal Dissociation Methods Differ for N—And O -Glycopeptides. Journal of Proteome Research, 19(8), 3286–3301. 10.1021/acs.jproteome.0c00218

24. Parker, B. L., Thaysen-Andersen, M., Fazakerley, D. J., Holliday, M., Packer, N. H., & James, D. E. (2016). Terminal Galactosylation and Sialylation Switching on Membrane Glycoproteins upon TNF-Alpha-Induced Insulin Resistance in Adipocytes. Molecular & Cellular Proteomics, 15(1), 141–153. 10.1074/mcp.M115.054221

25. Yu, Q., Wang, B., Chen, Z., Urabe, G., Glover, M. S., Shi, X., Guo, L.-W., Kent, K. C., & Li, L. (2017). Electron-Transfer/Higher-Energy Collision Dissociation (EThcD)-Enabled Intact Glycopeptide/Glycoproteome Characterization. Journal of the American Society for Mass Spectrometry, 28(9), 1751–1764. 10.1007/s13361-017-1701-4

26. Liu, M.-Q., Zeng, W.-F., Fang, P., Cao, W.-Q., Liu, C., Yan, G.-Q., Zhang, Y., Peng, C., Wu, J.-Q., Zhang, X.-J., Tu, H.-J., Chi, H., Sun, R.-X., Cao, Y., Dong, M.-Q., Jiang, B.-Y., Huang, J.-M., Shen, H.-L., Wong, C. C. L., … Yang, P.-Y. (2017). pGlyco 2.0 enables precision N-glycoproteomics with comprehensive quality control and one-step mass spectrometry for intact glycopeptide identification. Nature Communications, 8(1), 438. 10.1038/s41467-017-00535-2

27. Cheng, K., Chen, R., Seebun, D., Ye, M., Figeys, D., & Zou, H. (2014). Large-scale characterization of intact N-glycopeptides using an automated glycoproteomic method. Journal of Proteomics, 110, 145–154. 10.1016/j.jprot.2014.08.006

28. Sun, S., Shah, P., Eshghi, S. T., Yang, W., Trikannad, N., Yang, S., Chen, L., Aiyetan, P., Höti, N., Zhang, Z., Chan, D. W., & Zhang, H. (2016). Comprehensive analysis of protein glycosylation by solid-phase extraction of N-linked glycans and glycosite-containing peptides. Nature Biotechnology, 34(1), 84–88. 10.1038/nbt.3403

29. Dang, L., Shen, J., Zhao, T., Zhao, F., Jia, L., Zhu, B., Ma, C., Chen, D., Zhao, Y., & Sun, S. (2019). Recognition of Bisecting N -Glycans on Intact Glycopeptides by Two Characteristic Ions in Tandem Mass Spectra. Analytical Chemistry, 91(9), 5478–5482. 10.1021/acs.analchem.8b05639

30. Shah, P., Wang, X., Yang, W., Toghi Eshghi, S., Sun, S., Hoti, N., Chen, L., Yang, S., Pasay, J., Rubin, A., & Zhang, H. (2015). Integrated Proteomic and Glycoproteomic Analyses of Prostate Cancer Cells Reveal Glycoprotein Alteration in Protein Abundance and Glycosylation*. Molecular & Cellular Proteomics, 14(10), 2753–2763. 10.1074/mcp.M115.047928

31. Yang, W., Shah, P., Toghi Eshghi, S., Yang, S., Sun, S., Ao, M., Rubin, A., Jackson, J. B., & Zhang, H. (2014). Glycoform Analysis of Recombinant and Human Immunodeficiency Virus Envelope Protein gp120 via Higher Energy Collisional Dissociation and Spectral-Aligning Strategy. Analytical Chemistry, 86(14), 6959–6967. 10.1021/ac500876p

32. Lee, J. Y., Lee, H. K., Park, G. W., Hwang, H., Jeong, H. K., Yun, K. N., Ji, E. S., Kim, K. H., Kim, J. S., Kim, J. W., Yun, S. H., Choi, C.-W., Kim, S. I., Lim, J.-S., Jeong, S.-K., Paik, Y.-K., Lee, S.-Y., Park, J., Kim, S. Y., … Yoo, J. S. (2016). Characterization of Site-Specific N -Glycopeptide Isoforms of α-1-Acid Glycoprotein from an Interlaboratory Study Using LC–MS/MS. Journal of Proteome Research, 15(12), 4146–4164. 10.1021/acs.jproteome.5b01159

33. Yang, G., Hu, Y., Sun, S., Ouyang, C., Yang, W., Wang, Q., Betenbaugh, M., & Zhang, H. (2018). Comprehensive Glycoproteomic Analysis of Chinese Hamster Ovary Cells. Analytical Chemistry, 90(24), 14294–14302. 10.1021/acs.analchem.8b03520

34. Atashi, M., Reyes, C. D. G., Sandilya, V., Purba, W., Ahmadi, P., Hakim, Md. A., Kobeissy, F., Plazzi, G., Moresco, M., Lanuzza, B., Ferri, R., & Mechref, Y. (2023). LC-MS/MS Quantitation of HILIC-Enriched N-glycopeptides Derived from Low-Abundance Serum Glycoproteins in Patients with Narcolepsy Type 1. Biomolecules, 13(11), 1589. 10.3390/biom13111589

35. Watanabe, Y., Allen, J. D., Wrapp, D., McLellan, J. S., & Crispin, M. (2020). Site-specific glycan analysis of the SARS-CoV-2 spike. Science, 369(6501), 330–333. 10.1126/science.abb9983

36. Gashash, E. A., Aloor, A., Li, D., Zhu, H., Xu, X.-Q., Xiao, C., Zhang, J., Parameswaran, A., Song, J., Ma, C., Xiao, W., & Wang, P. G. (2017). An Insight into Glyco-Microheterogeneity of Plasma von Willebrand Factor by Mass Spectrometry. Journal of Proteome Research, 16(9), 3348–3362. 10.1021/acs.jproteome.7b00359

37. Pan, J., Hu, Y., Sun, S., Chen, L., Schnaubelt, M., Clark, D., Ao, M., Zhang, Z., Chan, D., Qian, J., & Zhang, H. (2020). Glycoproteomics-based signatures for tumor subtyping and clinical outcome prediction of high-grade serous ovarian cancer. Nature Communications, 11(1), 6139. 10.1038/s41467-020-19976-3

38. Carnielli, C. M., Melo De Lima Morais, T., Malta De Sá Patroni, F., Prado Ribeiro, A. C., Brandão, T. B., Sobroza, E., Matos, L. L., Kowalski, L. P., Paes Leme, A. F., Kawahara, R., & Thaysen-Andersen, M. (2023). Comprehensive Glycoprofiling of Oral Tumors Associates N-Glycosylation With Lymph Node Metastasis and Patient Survival. Molecular & Cellular Proteomics, 22(7), 100586. 10.1016/j.mcpro.2023.100586

39. Kawahara, R., Ugonotti, J., Chatterjee, S., Tjondro, H. C., Loke, I., Parker, B. L., Venkatakrishnan, V., Dieckmann, R., Sumer-Bayraktar, Z., Karlsson-Bengtsson, A., Bylund, J., & Thaysen-Andersen, M. (2023). Glycoproteome remodeling and organelle-specific N -glycosylation accompany neutrophil granulopoiesis. Proceedings of the National Academy of Sciences, 120(36), e2303867120. 10.1073/pnas.2303867120

40. Nayak, S., Zhao, Y., Mao, Y., & Li, N. (2021). System-Wide Quantitative N-Glycoproteomic Analysis from K562 Cells and Mouse Liver Tissues. Journal of Proteome Research, 20(11), 5196–5202. 10.1021/acs.jproteome.1c00451

41. Liu, S., Wang, H., Jiang, X., Ji, Y., Wang, Z., Zhang, Y., Wang, P., & Xiao, H. (2022). Integrated N - glycoproteomics Analysis of Human Saliva for Lung Cancer. Journal of Proteome Research, 21(7), 1589–1602. 10.1021/acs.jproteome.1c00701

42. Zhao, Y., Nayak, S., Raidas, S., Guo, L., Della Gatta, G., Koppolu, S., Halasz, G., Montasser, M. E., Shuldiner, A. R., Mao, Y., & Li, N. (2023). In-Depth Mass Spectrometry Analysis Reveals the Plasma Proteomic and N-Glycoproteomic Impact of an Amish-Enriched Cardioprotective Variant in B4GALT1. Molecular & Cellular Proteomics, 22(8), 100595. 10.1016/j.mcpro.2023.100595

43. Shu, Q., Li, M., Shu, L., An, Z., Wang, J., Lv, H., Yang, M., Cai, T., Hu, T., Fu, Y., & Yang, F. (2020). Large-scale Identification of N-linked Intact Glycopeptides in Human Serum using HILIC Enrichment and Spectral Library Search. Molecular & Cellular Proteomics, 19(4), 672–689. 10.1074/mcp.RA119.001791

44. Tian, W., Li, D., Zhang, N., Bai, G., Yuan, K., Xiao, H., Gao, F., Chen, Y., Wong, C. C. L., & Gao, G. F. (2021). O-glycosylation pattern of the SARS-CoV-2 spike protein reveals an “O-Follow-N” rule. Cell Research, 31(10), 1123–1125. 10.1038/s41422-021-00545-2

45. Hu, Z., Gao, W., Liu, R., Yang, J., Han, R., Li, J., Yu, J., Ma, D., & Tang, K. (2024). An efficient strategy with a synergistic effect of hydrophilic and electrostatic interactions for simultaneous enrichment of N— And O -glycopeptides. The Analyst, 149(4), 1090–1101. 10.1039/D3AN01888A

46. Escribano, J., Lopex-Otin, C., Hjerpe, A., Grubb, A., & Mendez, E. (1990). Location and characterization of the three carbohydrate prosthetic groups of human protein HC. FEBS Letters, 266(1–2), 167–170. 10.1016/0014-5793(90)81531-R

47. Chandrasekhar, K. D., Lvov, A., Terrenoire, C., Gao, G. Y., Kass, R. S., & Kobertz, W. R. (2011). O - glycosylation of the cardiac I Ks complex. The Journal of Physiology, 589(15), 3721–3730. 10.1113/jphysiol.2011.211284

48. Riethmueller, S., Somasundaram, P., Ehlers, J. C., Hung, C.-W., Flynn, C. M., Lokau, J., Agthe, M., Düsterhöft, S., Zhu, Y., Grötzinger, J., Lorenzen, I., Koudelka, T., Yamamoto, K., Pickhinke, U., Wichert, R., Becker-Pauly, C., Rädisch, M., Albrecht, A., Hessefort, M., … Garbers, C. (2017). Proteolytic Origin of the Soluble Human IL-6R In Vivo and a Decisive Role of N-Glycosylation. PLOS Biology, 15(1), e2000080. 10.1371/journal.pbio.2000080

49. Chien, Y.-C., Wang, Y.-S., Sridharan, D., Kuo, C.-W., Chien, C.-T., Uchihashi, T., Kato, K., Angata, T., Meng, T.-C., Hsu, S.-T. D., & Khoo, K.-H. (2023). High Density of N- and O-Glycosylation Shields and Defines the Structural Dynamics of the Intrinsically Disordered Ectodomain of Receptor-type Protein Tyrosine Phosphatase Alpha. JACS Au, 3(7), 1864–1875. 10.1021/jacsau.3c00124

50. Chernykh, A., Abrahams, J. L., Grant, O. C., Kambanis, L., Sumer-Bayraktar, Z., Ugonotti, J., Kawahara, R., Corcilius, L., Payne, R. J., Woods, R. J., & Thaysen-Andersen, M. (2024). Position-specific N- and O-glycosylation of the reactive center loop impacts neutrophil elastase–mediated proteolysis of corticosteroid-binding globulin. Journal of Biological Chemistry, 300(1), 105519. 10.1016/j.jbc.2023.105519

51. Rangel-Angarita, V., Mahoney, K. E., Kwon, C., Sarker, R., Lucas, T. M., & Malaker, S. A. (2023). False-Positive Glycopeptide Identification via In-FAIMS Fragmentation. JACS Au, 3(9), 2498–2509. 10.1021/jacsau.3c00264

52. Chongsaritsinsuk, J., Steigmeyer, A. D., Mahoney, K. E., Rosenfeld, M. A., Lucas, T. M., Smith, C. M., Li, A., Ince, D., Kearns, F. L., Battison, A. S., Hollenhorst, M. A., Judy Shon, D., Tiemeyer, K. H., Attah, V., Kwon, C., Bertozzi, C. R., Ferracane, M. J., Lemmon, M. A., Amaro, R. E., & Malaker, S. A. (2023). Glycoproteomic landscape and structural dynamics of TIM family immune checkpoints enabled by mucinase SmE. Nature Communications, 14(1), 6169. 10.1038/s41467-023-41756-y

53. Stavenhagen, K., Kayili, H. M., Holst, S., Koeleman, C. A. M., Engel, R., Wouters, D., Zeerleder, S., Salih, B., & Wuhrer, M. (2018). N- and O-glycosylation Analysis of Human C1-inhibitor Reveals Extensive Mucin-type O-Glycosylation. Molecular & Cellular Proteomics, 17(6), 1225–1238. 10.1074/mcp.RA117.000240

54. Larrucea, S., Butta, N., Arias-Salgado, E. G., Alonso-Martin, S., Ayuso, M. S., & Parrilla, R. (2008). Expression of podocalyxin enhances the adherence, migration, and intercellular communication of cells. Experimental Cell Research, 314(10), 2004–2015. 10.1016/j.yexcr.2008.03.009

55. Varki, A. (2001). Loss of N-glycolylneuraminic acid in humans: Mechanisms, consequences, and implications for hominid evolution. American Journal of Physical Anthropology, 116(S33), 54–69. 10.1002/ajpa.10018

56. Steentoft, C., Vakhrushev, S. Y., Joshi, H. J., Kong, Y., Vester-Christensen, M. B., Schjoldager, K. T.-B. G., Lavrsen, K., Dabelsteen, S., Pedersen, N. B., Marcos-Silva, L., Gupta, R., Paul Bennett, E., Mandel, U., Brunak, S., Wandall, H. H., Levery, S. B., & Clausen, H. (2013). Precision mapping of the human O-GalNAc glycoproteome through SimpleCell technology. The EMBO Journal, 32(10), 1478–1488. 10.1038/emboj.2013.79

57. Windwarder, M., & Altmann, F. (2014). Site-specific analysis of the O-glycosylation of bovine fetuin by electron-transfer dissociation mass spectrometry. Journal of Proteomics, 108, 258–268. 10.1016/j.jprot.2014.05.022

58. Solecka, B. A., Weise, C., Laffan, M. A., & Kannicht, C. (2016). Site-specific analysis of von Willebrand factor O-glycosylation. Journal of Thrombosis and Haemostasis, 14(4), 733–746. 10.1111/jth.13260

59. Cummings, R. D., Etzler, M., Hahn, M. G., Darvill, A., Godula, K., Woods, R. J., & Mahal, L. K. (2022). Glycan-Recognizing Probes as Tools. In A. Varki, R. D. Cummings, J. D. Esko, P. Stanley, G. W. Hart, M. Aebi, D. Mohnen, T. Kinoshita, N. H. Packer, J. H. Prestegard, R. L. Schnaar, & P. H. Seeberger (Eds.), Essentials of Glycobiology (4th ed.). Cold Spring Harbor Laboratory Press. http://www.ncbi.nlm.nih.gov/books/NBK579992/

60. Darula, Z., Sarnyai, F., & Medzihradszky, K. F. (2016). O-glycosylation sites identified from mucin core-1 type glycopeptides from human serum. Glycoconjugate Journal, 33(3), 435–445. 10.1007/s10719-015-9630-6

61. Malaker, S. A., Riley, N. M., Shon, D. J., Pedram, K., Krishnan, V., Dorigo, O., & Bertozzi, C. R. (2022). Revealing the human mucinome. Nature Communications, 13(1), 3542. 10.1038/s41467-022-31062-4

62. Mahoney, K. E., Chang, V., Lucas, T. M., Maruszko, K., & Malaker, S. A. (2024). Mass Spectrometry-Compatible Elution Technique Enables an Improved Mucin-Selective Enrichment Strategy to Probe the Mucinome. Analytical Chemistry, 96(13), 5242–5250. 10.1021/acs.analchem.3c05762

63. Kufe, D. W. (2009). Mucins in cancer: Function, prognosis and therapy. Nature Reviews Cancer, 9(12), 874–885. 10.1038/nrc2761

64. Bakkers, M. J. G., Moon-Walker, A., Herlo, R., Brusic, V., Stubbs, S. H., Hastie, K. M., Saphire, E. O., Kirchhausen, T. L., & Whelan, S. P. J. (2022). CD164 is a host factor for lymphocytic choriomeningitis virus entry. Proceedings of the National Academy of Sciences, 119(10), e2119676119. 10.1073/pnas.2119676119

65. Pinho, S. S., & Reis, C. A. (2015). Glycosylation in cancer: Mechanisms and clinical implications. Nature Reviews Cancer, 15(9), 540–555. 10.1038/nrc3982

66. Hellman, N. E., & Gitlin, J. D. (2002). CERULOPLASMIN METABOLISM AND FUNCTION. Annual Review of Nutrition, 22(1), 439–458. 10.1146/annurev.nutr.22.012502.114457

67. Loo, T., Patchett, M.L., Norris, G.E., & Lott, J.S. (2002). Using secretion to solve a solubility problem: high-yield expression in Escherichia coli and purification of the bacterial glycoamidase PNGase F. Protein Expr. Purif., 24(1), 90–98. 10.1006/prep.2001.1555.

