## Supplemental Information for "Quantification and site-specific analysis of co-occupied N- and O-glycopeptides"

**This PDF file includes:**

Figures S1 to S3

**Additional supplementary materials not included in this document:**

Supplemental Table 1 – Previously identified co-occupied glycopeptides in the literature
Supplemental Table 2 – SmE, trypsin, and PNGaseF digest of TIM-4 output and manually validated peptides
Supplemental Table 3 – Calculated variables for co-occupancy
Supplemental Table 4 – SmE, trypsin, and PNGaseF digest of C1-Inh output and manually validated peptides
Supplemental Table 5 – StcE, trypsin, and PNGaseF digest of podocalyxin output and manually validated peptides
Supplemental Table 6 – IMPa, trypsin, and PNGaseF digest of podocalyxin output and manually validated peptides
Supplemental Table 7 – SmE, trypsin, and PNGaseF digest of ConA-enriched HeLa lysate output and manually validated peptides
Supplemental Table 8 – IMPa, trypsin, and PNGaseF digest of Jacalin-enriched human serum output and manually validated peptides
Supplemental Table 9 – SmE and PNGaseF digest of StcE-enriched HeLa lysate output and manually validated peptides

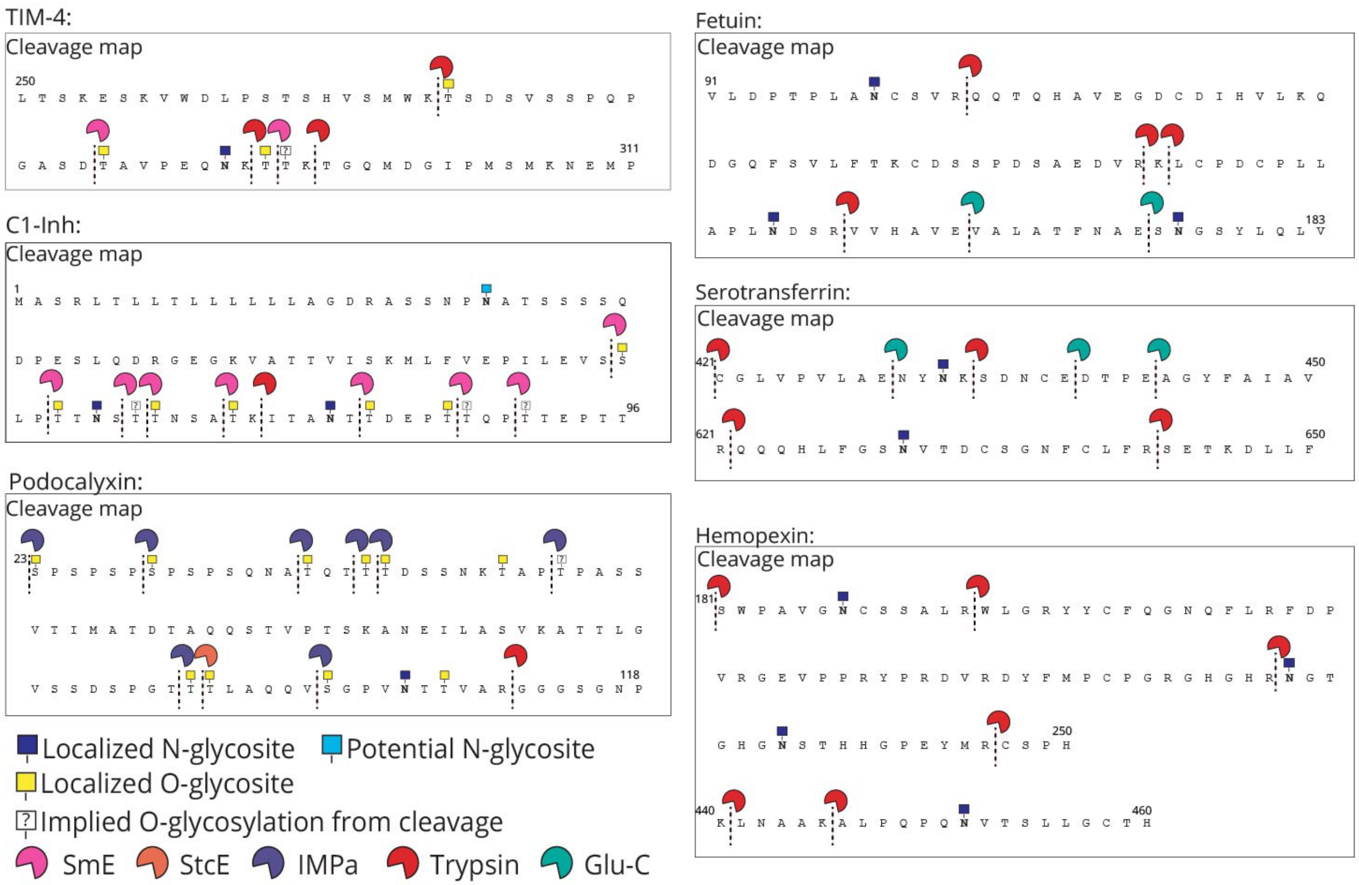


**Supplementary Figure 1. Map of all cleavage events.** Proteins were digested with a combination of proteases including SmE (pink), StcE (orange), ImpA (blue), trypsin (red), and/or GluC (green), followed by MS analysis and manual data interpretation. All cleavage events around N-sequons in each glycoprotein (mucins and non-mucin glycoproteins). Cleavage events are denoted by dashed lines with the appropriate protease. Implied sites from mucinase cleavages are denoted with white boxes.

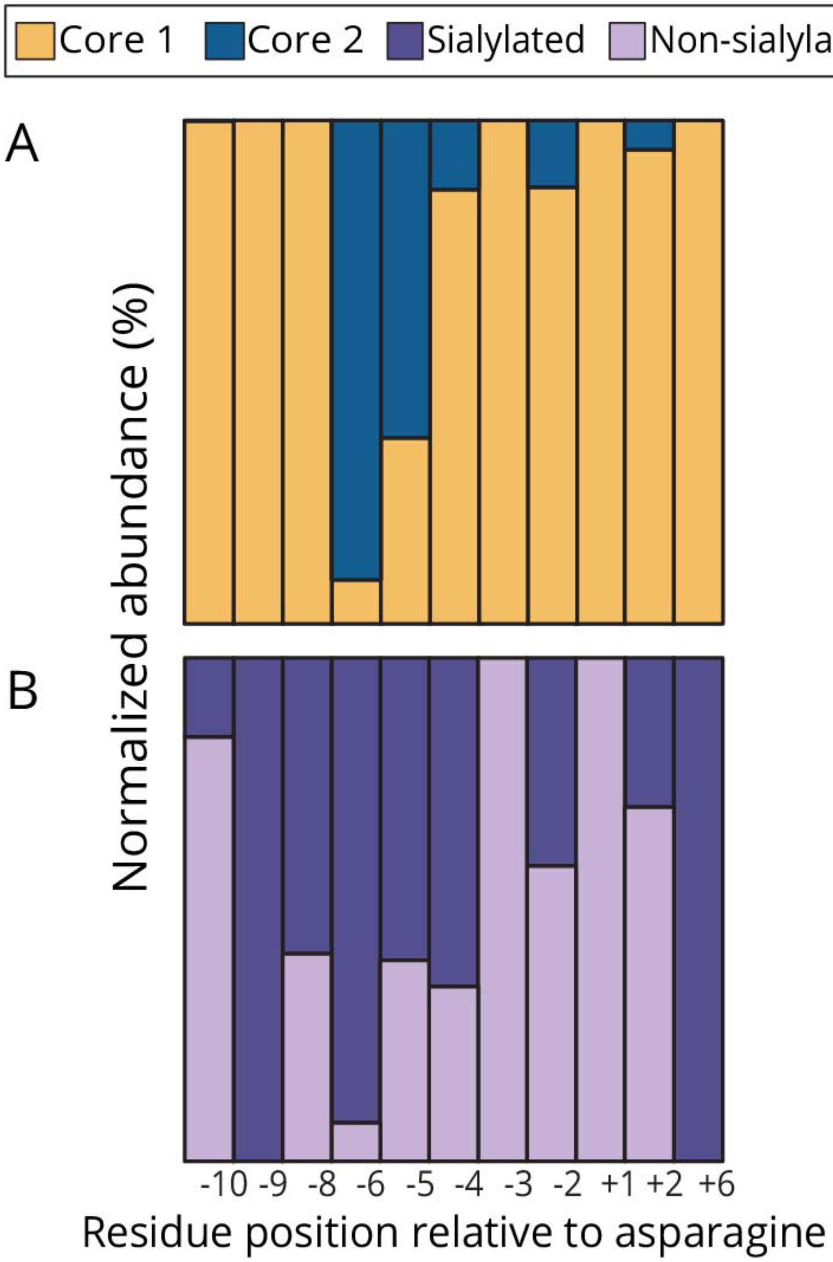


**Supplementary Figure 2. Analysis of co-occupied O-glycosite compositions in mucin-domain glycoproteins.** Four mucins were digested with a combination of proteases and subjected to MS analysis followed by manual data curation. Using Thermo XCalibur, extracted ion chromatograms were generated for all identified glycopeptides. We normalized residue positions within peptides to the occupied Asn. Residues N-terminal to the Asn are considered negative while residues C-terminal to the Asn are considered positive. The AUCs of each glycopeptide containing the corresponding glycan were calculated then summed up to determine relative abundance of various glycoforms for each normalized position.

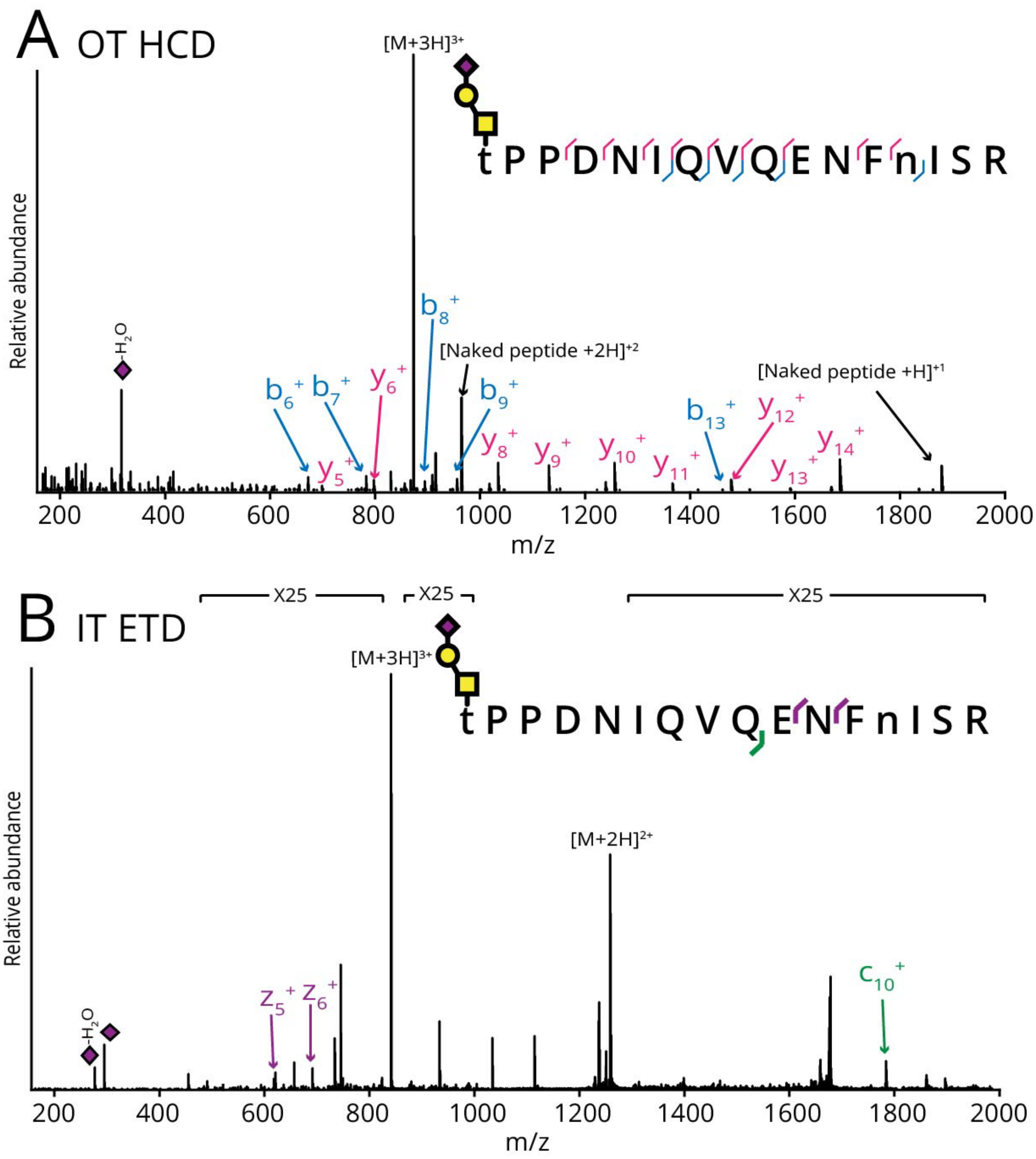


**Supplementary Figure 3. Co-occupancy of Thr24 and Asn36 in AMBP.** Human serum was subjected to digestion with ImpA, trypsin, and PNGaseF followed by Jacalin enrichment, MS analysis, and manual data analysis. A) HCD spectrum and B) ETD spectrum of a co-occupied glycopeptide from AMBP. The spectra allow for unambiguous localization of H1N1A1 to Thr24, as well as deamidation of Asn36, indicating N-glycosylation.
